# Impact of Training Regimes and Task Similarity on Learning and Cognitive Transfer

**DOI:** 10.1101/2025.09.22.677779

**Authors:** N. Menghi, S. Vigano’, W.J. Johnston, S. Elnagar, S. Fusi, F.C. Doeller

**Affiliations:** Max Planck Institute for Human Cognitive and Brain Sciences, Department of Psychology, Germany; Center for Mind/Brain Sciences, University of Trento, Rovereto 38068, Italy; Center for Theoretical Neuroscience Columbia University, New York, NY, USA, Mortimer B. Zuckerman Mind, Brain and Behavior Institute, Columbia University, New York, NY, USA; Kavli Institute for Systems Neuroscience, Center for Neural Computation, Jebsen Center for Alzheimer’s Disease, Norwegian University of Science and Technology, Trondheim 7491, Norway

**Keywords:** Continual Learning, Transfer Learning, Interference, Training Regime, Curriculum Learning

## Abstract

Learning depends not only on the content of what we learn, but also on how we learn and on how experiences are structured over time. To investigate how task similarity and training regime interact during learning, we trained participants on spatial and conceptual learning tasks that shared either similar or distinct underlying structures, using either interleaved or blocked regimes. Interleaving the two tasks hindered performance when their structures were similar, compared to when they were different. In contrast, blocked training produced the opposite effect: it improved performance and facilitated transfer across similar tasks. This effect, however, emerged only when participants first learned the conceptual task, followed by the spatial task, suggesting an asymmetric interaction between task order and structural similarity. We also replicated our results using a neural network model, providing converging evidence for the computational principles governing the interplay between training regime and structural similarity in multi-task learning.

## 1 Introduction

Unlike machines, humans exhibit an extraordinary capacity for learning and adapting to a wide range of tasks, demonstrating resilience against interference and forgetting. We can effortlessly learn to navigate through vast cities in the United States while retaining our spatial memory and orientation of smaller, irregular towns in Italy. Similarly, we can acquire new languages without forgetting our native tongue. Unraveling the principles of how “continual learning” (Parisi, Kemker, Part, Kanan, & Wermter, 2019; Flesch, Balaguer, Dekker, Nili, & Summerfield, 2018; Hadsell, Rao, Rusu, & Pascanu, 2020; Kirkpatrick et al., 2017; Fusi, 2021; Flesch, Saxe, & Summerfield, 2023) works is crucial not only for comprehending our own cognitive processes but also for guiding contemporary artificial intelligence research, especially in the face of the “catastrophic forgetting” artificial neural networks suffer from. Despite its significance, however, the precise mechanisms responsible for driving this ability remain unresolved (Kudithipudi et al., 2022).

A growing body of work suggests that understanding continual learning requires understanding how the brain organizes knowledge and the structure of experiences. According to this perspective, the brain extracts structural regularities from experience and uses them to build internal models of the world that support generalization, prediction, and decision-making (Behrens et al., 2018; Bellmund, Gärdenfors, Moser, & Doeller, 2018; Kaplan, Schuck, & Doeller, 2017; Whittington et al., 2020).

Within this framework, items, concepts, or “states” can be organized in relational maps where their relative positions are determined by a set of relevant features, dimensions, or outcomes (Shepard & Metzler, 1971; Gardenfors, 2004; Bellmund et al., 2018), and distances in these maps code for similarity, placing similar inputs closer than dissimilar ones (Viganò, Rubino, Di Soccio, Buiatti, & Piazza, 2021; Viganò, Bayramova, Doeller, & Bottini, 2023; Theves, Fernandez, & Doeller, 2019; Theves, Fernández, & Doeller, 2020; Nitsch, Garvert, Bellmund, Schuck, & Doeller, 2024; Barnaveli, Viganò, Reznik, Haggard, & Doeller, 2025). Extracting these models not only facilitates storage and memorization but also promotes generalization and the transfer of acquired knowledge to similar situations (Mark, Moran, Parr, Kennerley, & Behrens, 2020; Aho, Roads, & Love, 2022; Dekker, Otto, & Summerfield, 2022; Zhang & Yu, 2024; Huang & Doeller, 2026). For example, after learning the water and light requirements for one plant in our garden, this information can be easily applied to new exemplars or species that share sufficient similarities, avoiding the need to (re)learn from scratch. Inspired also by intuitions from machine learning (e.g., (Caruana, 1997)), prior studies have indeed demonstrated that humans exploit structural similarities between tasks, and that this can enhance knowledge transfer and generalization (Menghi, Kacar, & Penny, 2021; Wu, Schulz, Garvert, Meder, & Schuck, 2020; K. Garner, Lynch, & Dux, 2016; Holton, Braun, Thompson, Grohn, & Summerfield, 2025).

Directly comparing humans and machines, Flesch and colleagues (2018) observed that humans, but not machines, benefit from learning different, orthogonal, classification tasks in separated and temporally autocorrelated contexts. This blocked training regime prevented interference between tasks even on later generalization tests, demonstrating humans’ ability to decompose complex tasks into non-interfering subcomponents. According to their perspective, this approach might support continual learning by facilitating participants to create separated and non-overlapping low-dimensional embeddings of the stimulus space, which thus align with the behaviorally relevant orthogonal rules. In other words, a blocked training regime helps to differentiate what needs to be differentiated, efficiently meeting the requirements of a “continual learning” system. Following this logic, one might interpret the factorization/separation proposed by Flesch and colleagues as a mechanism employed by the brain to specifically protect against the interference that very different tasks might produce. In other words, to ensure that the system is not overloaded by contrasting information, contextual dissociation proves beneficial, making a blocked regime favorable for decorrelation/differentiation compared to an interleaved one. If this is the case, then the effect should reverse when the subspaces are not orthogonal but rather similar. In such instances, the system should have better performance when presented with multiple tasks of similar structure in an interleaved fashion, or at least it should remain unchanged: we might get confused if, in our garden, we constantly alternate taking care of plants with very different light or water requirements, but if they all share the same needs, our cost should be highly reduced.

In a previous study, we tested the above hypothesis by designing an experiment with two tasks from different domains — spatial and conceptual — each featuring either similar or distinct structures, under an interleaved training regime. Contrary to our expectations, the interleaved regime did not facilitate transfer; instead, it caused significant interference between tasks with similar structures during learning (Menghi, Johnston, et al., 2025). These results highlighted a more complex interplay between task similarities and training regime than previously assumed. To further characterize this interplay, in the current study, we test both human participants and a neural network model under a blocked training regime, and compare the results to those we previously obtained under an interleaved training regime. Specifically, we reuse the same neural network architecture and task design from our prior study (Menghi, Johnston, et al., 2025), but modify the training regime to present tasks in separate blocks rather than in an interleaved fashion. Crucially, our goal is not only to examine the individual effects of task structure similarity and training regime, but to understand how these two factors interact to shape learning dynamics. While prior work has often treated task similarity and training regime as independent variables, their interaction may be key to understanding when generalization is facilitated and when interference occurs. Investigating this interaction will provide insight into the conditions under which shared structure is beneficial versus detrimental, offering a more comprehensive account of how humans and machines build internal representations for continual learning. We predict that participants trained under a blocked regime will perform better during the learning phase compared to those trained interleaved, although this advantage may not persist during the subsequent testing phase (see also (Flesch et al., 2018)). Furthermore, we expect both the neural network and participants to transfer knowledge between tasks during blocked training when structures are shared.

In our behavioral data, we find evidence of transfer between tasks under blocked training. Interestingly, and in contrast to previous findings (Wu, Schulz, Garvert, Meder, & Schuck, 2018; Wu et al., 2020; Lerousseau & Summerfield, 2024), this transfer appears to be directional, from conceptual to spatial tasks. We discuss the reason for this unexpected asymmetry in the Discussion. Consistent with our predictions, the neural network simulations also exhibit transfer between blocked consecutive tasks, but only when the tasks share the same underlying structure, reinforcing the idea that structural similarity facilitates generalization under specific training regimes.

## 2 Results

We trained participants to perform an associative learning task across two contexts in either an interleaved regime or a blocked one; data for the interleaved training regime were collected as part of a different experiment (see (Menghi, Johnston, et al., 2025)). They had to predict if a seed was going to “grow” or “die” based on two distinct sets of features, where the relevant features were determined by the identity of the seed (See Fig. 1A-B). Decisions about the first seed relied on what we called the “conceptual features”: participants had to learn the association between configurations of the opacity of a sun and a drop of water images and the outcome (grow or die). Decisions about the second seed relied on “spatial features”: participants had to learn the association between configurations of position in space in which the seed was planted and an outcome (grow or die). The different contexts were cued by two different seeds (See Fig. 1A).

**Figure 1:**
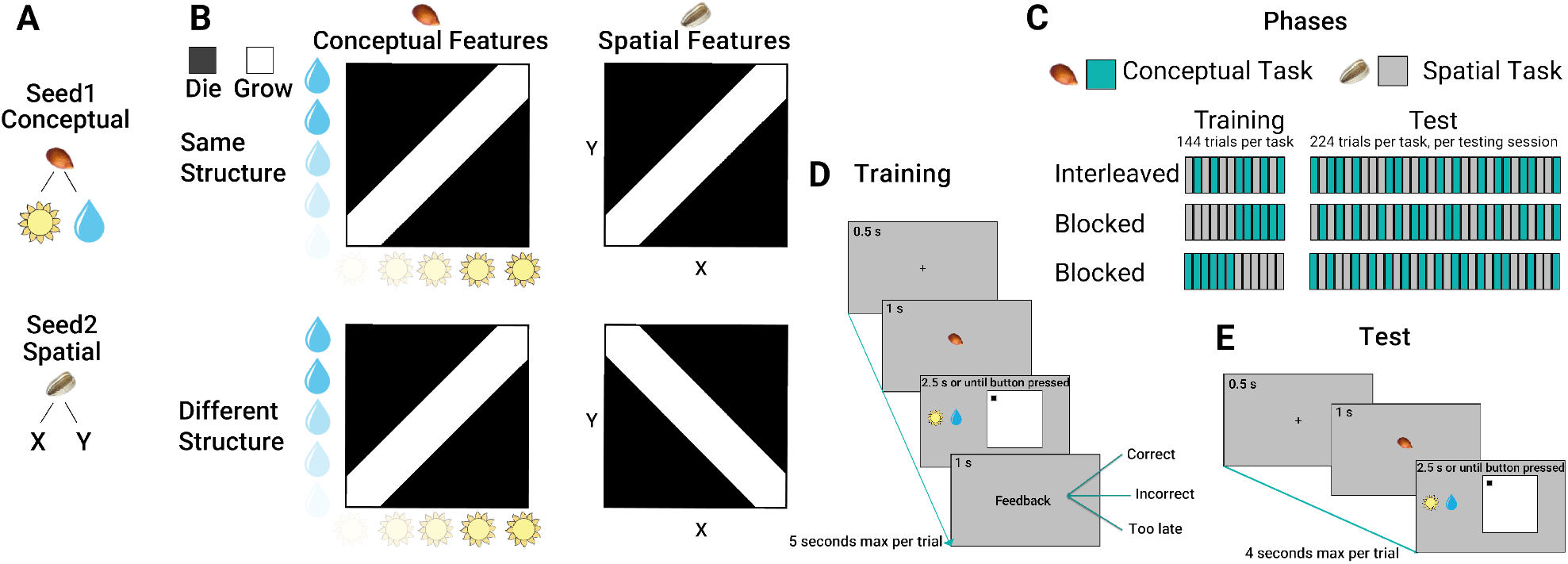
Stimuli, Task Structures and Trial Structures. **Panel A** shows the experimental stimuli and cues. Two seeds were used to cue for the two different tasks, conceptual and spatial. In the conceptual task, we used a sun and a water drop symbol as features composing a stimulus configuration. In the spatial task, we used a position (x,y) on a square map as a stimulus. **Panel B** shows the different structures used. In the conceptual task, each vertical (sun opacity) and horizontal (a drop of water opacity) position can be combined, creating a continuous map, or structure, from which we draw stimuli. In these structures, black colour is associated with the outcome “die” and white is associated with the outcome “grow”. The structures of the two tasks were the same in one group and orthogonal in the other group of participants. **Panel C** shows the different phases of the experiment. In each phase, conceptual and spatial tasks were organized in either an interleaved or a blocked regime. Blocked regimes were further divided into two groups, one starting with the conceptual task, the other with the spatial one. **Panel D** shows the schematic of a trial structure during the training phase. A fixation cross was shown for 0.5 seconds, and then the seed cueing the task domain appeared for 1 second. Brown seed cued participants to pay attention to the conceptual features and ignore the spatial ones, and vice-versa, the grey seed cued participants to pay attention to the spatial features and ignore the conceptual ones. Afterwards, both spatial and conceptual configurations appeared on screen and stayed there for 2.5 seconds maximum, or until response. This is when participants used a button box to predict if, based on the relevant features presented, the seed was associated with “grow” or “die”. Finally, feedback appeared and stayed on screen for 1 second. **Panel E** shows the schematic of a trial structure during the testing phase. A fixation cross was shown for 0.5 seconds, and then the seed cueing the task domain appeared for 1 second. Afterwards, spatial and conceptual configurations appeared on screen and stayed there for 2.5 seconds maximum, or until response. Unlike the training phase, no feedback was provided.

Participants were randomly assigned to different groups, see table 1 in the methods section for a breakdown of the groups and participants assignments. Half of the participants were assigned to either a “Same structure” (SaSt) or a “Different structure” (DiSt) group, which differed based on the similarity between the feature to outcome mappings (see Fig.1B). Participants in the SaSt group had the same feature-outcome structure in both contexts, while participants in the DiSt group had a flipped structure across the two contexts (Fig.1B). 60 participants were assigned to the interleaved condition and 120 to the blocked condition. Participants in the interleaved group were trained and tested with an interleaved regime: trials from the conceptual and spatial task were sampled randomly (see Fig.1C). Participants in the blocked group were trained with a blocked regime: 144 trials in one task followed by 144 trials in the other one, followed by a testing session with an interleaved regime. Finally, we divided participants in the blocked condition into two groups, one starting with the spatial task, and one beginning with the conceptual one (see Table 1 for a breakdown of the experimental design). Each participant was first trained via feedback learning (see Fig.1D), and then tested (see Fig.1E) during two separate sessions, on the same day, separated by a navigation task that pertained to a different question and will be analysed separately (See Methods).

**Table 1:** Subjects and groups. Subjects were assigned to either the SaSt/DiSt or to blocked/Interleaved conditions. Numbers between brackets are the number of participants discarded.

|  | Interleaved | Blocked Spatial First | Blocked Conceptual First |
| --- | --- | --- | --- |
| SaSt | 30(3) | 30(1) | 30 |
| DiSt | 30(3) | 30(1) | 30(2) |

To assess the effect of task similarity and training regime on representation learning, we divided the analysis into three parts. First, we compared participant accuracies between the interleaved and the blocked group by assessing their learning and generalization performance, where generalization refers to performance on novel stimulus configurations not seen during training. Second, we looked at the effect of task similarity and its interaction with the training regime. Finally, we checked if there was a directionality of the effect by looking at the learning performance between participants who either started with the spatial or the conceptual tasks in the blocked training regime.

### 2.1 Training Regime

We operationalized learning accuracy as the proportion of correct responses over trials separately in the training and test phases. The test phase included both old and new stimuli. We then quantified generalization during the test session by computing the proportion of correct responses for novel configurations that, contrary to old ones, were not presented during training. To test differences across training regimes and phases, we conducted a mixed-design ANOVA with Group (blocked vs. interleaved; between-subjects factor) and Phase (training vs. test; within-subjects factor). Importantly, the mixed ANOVA revealed a significant main effect of Group (*F* (1, 176) = 18.038, *p* < 0.001, 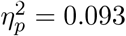), a significant main effect of Phase (*F* (1, 176) = 72.547, *p* < 0.001, 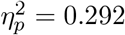), and a significant Group × Phase interaction (*F* (1, 176) = 14.312, *p* < 0.001, 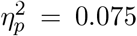), indicating that the performance difference between training regimes differed across training and test phases. We then performed two post hoc between-subject t-tests to further characterize the direction of the group differences during the training and test phases. We found that participants belonging to the blocked group performed better compared to those in the interleaved regime during both the training phase (t(168) = 6.693, p < 0.001, r = 0.459) and test phase (t(168) = 3.488, p < 0.001, r = 0.260), as shown in Figure 2 A and B. To look at the difference in performance between old and new stimuli, during the testing phase, we performed a 2×2 mixed-design ANOVA, with training regime as between factor and novelty of stimuli (new or old) as within factor. We found both main effects and the interaction to be statistically significant (*Regime* : *F* (1, 168) = 14.109, *p* < 0.001, 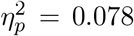, *Novelty* : *F* (1, 168) = 204.16, *p* < 0.001, 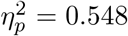, *Interaction* : *F* (1, 168) = 16.395, *p* < 0.001, 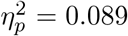). This indicates that, overall, participants performed better for the old stimuli compared to the new, unseen ones and that they had a better performance when trained with a blocked regime, but that this group difference was mostly led by their performance for the old stimuli, as shown in Figure 2 C. Indeed, despite being numerically different, the performance of the two groups for the new configurations during the test phase did not diverge in a statistically significant way (*p* = 0.113), suggesting that generalization happened under both training regimes, as both groups performed above chance (Blocked: *p* < 0.001; Interleaved: *p* < 0.001) See also Supplementary Figure 2 for a further illustration of this pattern.

**Figure 2:**
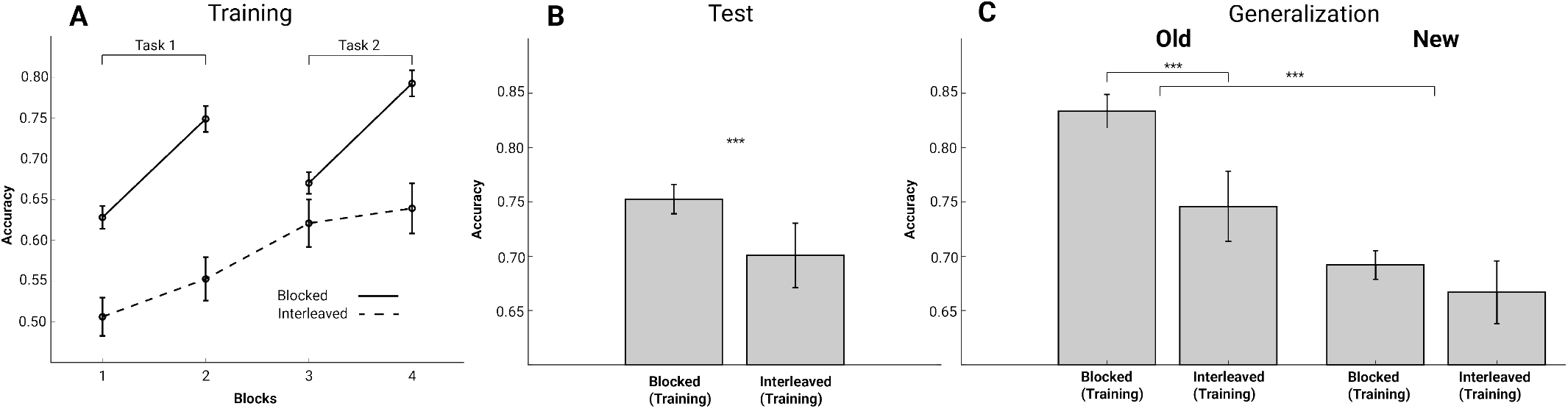
General performance and Generalization. **Panel A** shows participants’ mean accuracies during the training blocks, with each block consisting of 72 trials and **Panel B** during test phases. **Panel C** shows participants’ performance for old and new stimuli during the testing phase. Participants are divided into the different training regimes; note that the test phase was always interleaved across all participants. All the error bars show standard error of the mean.

### 2.2 Task Similarity and Training Regime

To directly examine whether task similarity interacted with training regime, we conducted two two-way ANOVAs on training and test performance with Regime (blocked vs. interleaved) and Structure (same vs. different) as between-subjects factors. During the training phase, this analysis revealed a significant main effect of Regime (*F* (1, 166) = 47.09, *p* < 0.001,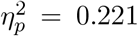), a significant main effect of Structure (*F* (1, 166) = 3.95, *p* = 0.049, 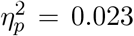), and a significant Regime *×* Structure interaction (*F* (1, 166) = 6.07, *p* = 0.015, 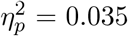). During the testing phase, the analysis revealed a significant main effect of Regime (*F* (1, 166) = 12.18, *p* < 0.001, 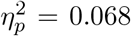), but neither the main effect of Structure (*F* (1, 166) = 0.77, *p* = 0.381, 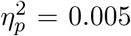) nor the Regime *×* Structure interaction reached significance (*F* (1, 166) = 2.75, *p* = 0.099, 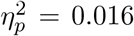). Together, these results suggest that the influence of task similarity was more pronounced during training than during testing. To further explore the interaction between training regime and task similarity in more detail, we ran separate analyses for blocked and interleaved regimes, looking at the difference in performance between SaSt and DiSt groups.

#### 2.2.1 Interleaved

We performed two between-subject t-tests to assess whether general performance differed between the task similarity groups in the interleaved training regimes. We found that participants belonging to the DiSt group performed better compared to those in the SaSt group during the training phase (t(52) = 2.908, p = 0.005, r = 0.374), as shown in Figure 3 A. Their performance, however, was not statistically different during the test phase (t(52) = 1.426, p = 0.159, r = 0.19). We then performed two 2×2 mixed-design ANOVAs, with between factor the group and within factor the novelty of the stimuli in the test phase. We found a significant main effect of novelty (*F* (1, 52) = 37.141, *p*<0.001, 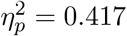) and interaction (*F* (1, 52) = 5.240, *p* = 0.026, 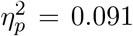), indicating that participants performed better with old stimuli compared to new ones, but no group main effect (*F* (1, 52) = 2.497, *p* = 0.12, 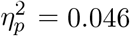) (See Figure 3C).

**Figure 3:**
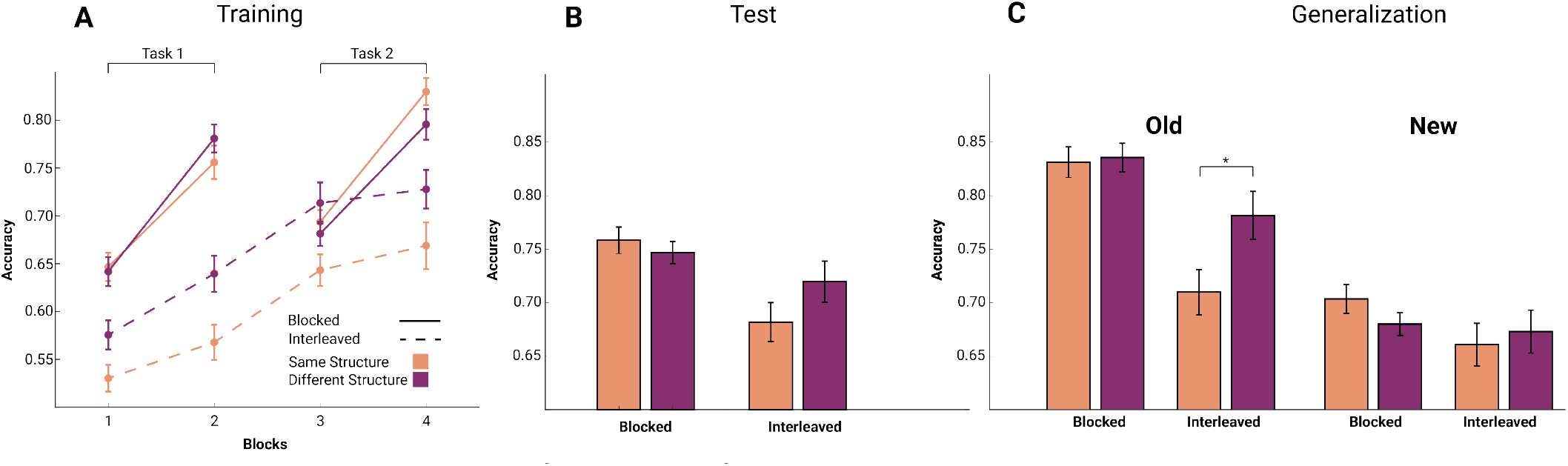
General performance and Generalization for SaSt and DiSt groups. **Panel A** shows participants’ mean accuracies during the training blocks and **Panel B** during test phases. Participants are divided into SaSt and DiSt group conditions and into the different training regimes groups. **Panel C** shows participants’ performance for old and new stimuli during the testing phase. Participants are divided into the different training regimes and into the different training regimes groups. All the error bars show the standard error of the mean.

#### 2.2.2 Blocked

We performed two between-subject t-tests to assess whether general performance differed between the task similarity groups in the blocked training regimes. We found no differences in performance between DiSt group and the SaSt group during the training phase (t(114) = 0.387, p = 0.699, , r = 0.04), nor during the testing phase (t(114) = 0.387, p = 0.699, r = 0.04) as shown in Figure 3 A and B. We then performed two 2×2 mixed-design ANOVAs, with between factor being the group (DiSt Vs SaSt) and within factor the novelty of the stimuli on the test phase. We found only the novelty main effect (*F* (1, 114) = 284.9, *p*<0.001, 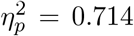) but no significant group effect nor interaction (*TaskSimilarityF* (1, 114) = 0.346, *p*=0.557, 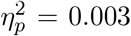 *InteractionF* (1, 114) = 2.766, *p* = 0.099, 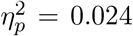), indicating that participants performed better with old stimuli compared to new ones (See Figure 3C).

### 2.3 Directionality of transfer learning in blocked regime

Next, we focused our analysis on the blocked group to see whether there is a directionality in the way task similarity influenced transfer during the training phase, specifically when switching from one task to the next. One possibility is that transfer is stronger when switching from a conceptual task to a spatial one, indicating that conceptual representations may support performance in more concrete spatial contexts. Conversely, it could be that transfer is more effective in the opposite direction—from spatial to conceptual—indicating that grounding in (more concrete) spatial representations may scaffold the acquisition of (more abstract) conceptual ones. For the spatial-first group, we compared performance on the conceptual task between the SaSt and DiSt conditions. Similarly, for the conceptual-first group, we compared performance on the spatial task between the SaSt and DiSt conditions. We performed a mixed-design ANOVA with Order (spatial-first vs. conceptual-first; between-subjects factor) and Structure (same vs. different; within-subjects factor). The ANOVA did not reveal significant main effect of Structure (*F* (1, 112) = 1.68, *p* = 0.1971, 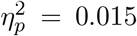) while Order (*F* (1, 112) = 2.99, *p* = 0.0867, 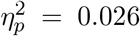) and order by structure interaction (*F* (1, 112) = 3.15, *p* = 0.079, 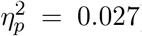) showed a trend-level effect. To further explore the pattern suggested by the interaction, we conducted exploratory between-subject t-tests and found that participants in the SaSt performed significantly better in the spatial task for participants starting with the conceptual one (t(56) = 2.162, p = 0.034, r = 0.28), but no differences in the conceptual task for participants starting with the spatial one (t(56) = 0.352, p = 0.726, r = 0.05) as shown in Figure **??**. The direction of transfer appears to be asymmetric, with a significant effect when transferring from conceptual to spatial tasks, but not the other way around.

### 2.4 Artificial neural networks offer insight into the difference in learning performance

To investigate how different training regimes influence learning under shared versus distinct task structures, we employed the same neural network architecture used in our previous work(Menghi, Johnston, et al., 2025). In the current study, however, the tasks were presented in a blocked fashion rather than interleaved as in the previous study. The networks received input variables specific to each task, along with additional inputs that signaled the context of each trial—analogous to the seed identity cues used in the behavioral experiment (see Methods). Some networks were trained on sequential tasks that shared the same underlying structure across, while others were trained on sequential tasks with different structures. We then assessed the learning speed for both the first and second tasks in the sequence.

We first tested the network using input variables assigned to perfectly orthogonal dimensions. We observed no difference in learning speed between networks trained on same-structure and different-structure tasks, for both the first and second tasks (Figure 5, panel D, left). This is because, from the network’s perspective, same- and different-structure conditions are indistinguishable: the orthogonality of the input variables prevents any shared representational overlap between the two task contexts, eliminating any potential for transfer or interference. To introduce a representational bias in the network, we manipulated the similarity between task features by representing the different features in the input along semi-orthogonal dimensions (Figure 5, panel C, middle), with similarity quantified using the cosine similarity between the vectors defining each feature. This type of representational geometry could be useful for making analogies between features that share a common similarity. For instance, it could serve as a representational substrate for analogizing more sun with being “further” along the x-axis. The degree to which these different features are viewed as similar to each other could be encoded in the degree of overlap between their feature dimensions in the neural population geometry.

Networks trained on tasks with higher feature similarity and the same task structure across contexts exhibited faster learning for the second task (Figure 5, panel E, middle). This pattern qualitatively mirrors the behavioral effect observed in our participants.

Finally, we examined an extreme version of this similarity manipulation in which the network received only two task-relevant features, perfectly aligned across contexts (Figure 5, panel C, right). In this scenario, the same-structure task becomes non-contextual: the network—or, by analogy, the participant—can ignore the contextual variable (e.g., seed identity) and still perform the task accurately. In contrast, for the different-structure condition, context remains essential for correct task execution. As a result, networks trained on the same-structure task learned the second task substantially faster than those trained on different-structure tasks (Figure 5, panel E, right).

Overall, the neural networks revealed that increasing similarity between relevant features enhances learning in tasks with the same underlying structure, which is consistent with our behavioral results. Interestingly, in the intermediate feature similarity regime (Figure 5, panels C, D, E, middle), this pattern depends on whether the network was trained in a block or interleaved manner. In particular, blocked training produces faster learning in the same structure condition, while interleaved training produces faster learning in the different structure condition – just as in our human participants.

## 3 Discussion

We investigated how training regime, task similarity, and their interaction influence representation learning. Participants learned two tasks under different training regimes, where stimulus–outcome associations were defined by a low-dimensional manifold that was either shared or distinct across tasks (see Fig. 1).

Our behavioral analyses revealed that a blocked training regime benefited learning during the training phase (see Fig. 2). Although test performance was also higher in the blocked condition, participants performed better on previously encountered configurations than on novel ones, indicating that this advantage was led by improved memorization rather than enhanced generalization. Supporting this, performance on novel stimuli did not differ between training regimes, suggesting no difference in generalization ability.

During training, participants exposed to interleaved regime with similar structures showed worse performance compared to participants trained with tasks with different structures, consistent with interference from overlapping representations (see (Menghi, Johnston, et al., 2025)). Importantly, this interference effect diminished over time, and group differences disappeared during the testing phase (see Fig. 3). Additionally, we observed a possible directional transfer effect in the blocked condition. Participants who began with the conceptual task and trained on structurally similar tasks performed better on the second, spatial, task compared to those trained on dissimilar structures (see Fig. 4). However, no such benefit was observed when participants started with the spatial task. This suggests a potential asymmetry in transfer from conceptual to spatial domains; however, this interpretation should be considered cautiously, given the lack of a significant interaction in the full model analysis.

**Figure 4:**
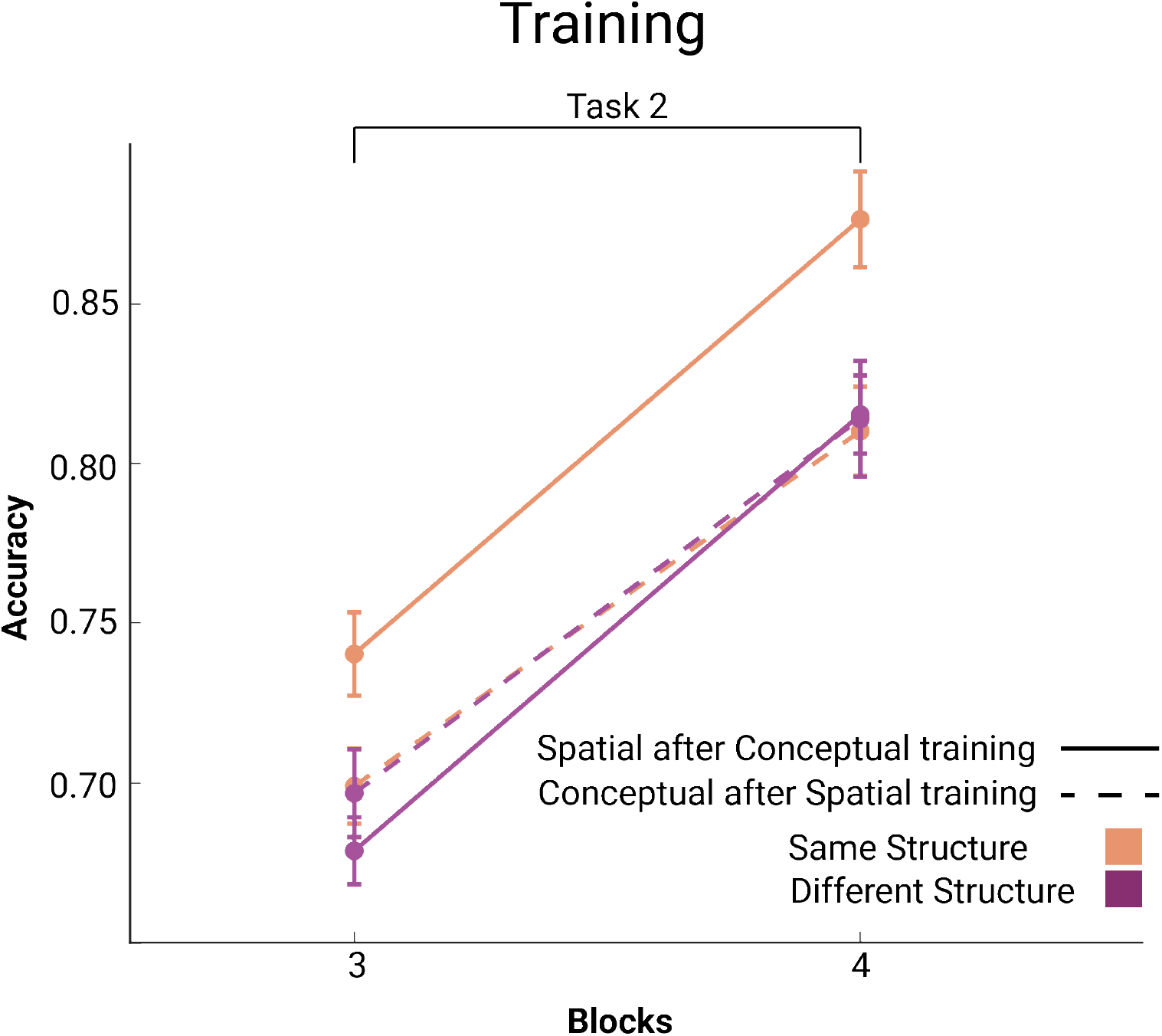
Transfer benefit during training. The figure shows participants’ mean accuracies, during blocked training regime, for the second task, which appears in the third and fourth blocks of the study. Participants are divided into SaSt and DiSt groups. Error bars show the standard error of the mean.

These results highlight the critical role of both training regime and task structure in shaping representation learning, demonstrating how blocked training can facilitate transfer when task structures align, while interleaved training may lead to interference, particularly when tasks share similarities.

The neural network model offered additional insight into the nature of the shared representations (see Fig. 5). When the two tasks involved orthogonal dimensions, task structure had no impact on the network’s performance—the same and different conditions were functionally indistinguishable from the network’s perspective. In contrast, as the similarity between task dimensions increased, the network learned more efficiently when the task structures were aligned, mirroring the effect observed in human participants. This pattern differed under an interleaved training regime, where the network experienced interference due to overlapping representations, consistent with our previous findings (Menghi, Johnston, et al., 2025). These results highlight an interaction between the training regime and task structure. Our human and network findings align with Holton et al. (Holton et al., 2025), who showed that both biological and artificial learners benefit from shared representations during learning. However, they also revealed the flip side of this effect: reusing existing representations can lead to interference with previously acquired knowledge (See also (K. G. Garner & Dux, 2023; Musslick & Cohen, 2021).

**Figure 5:**
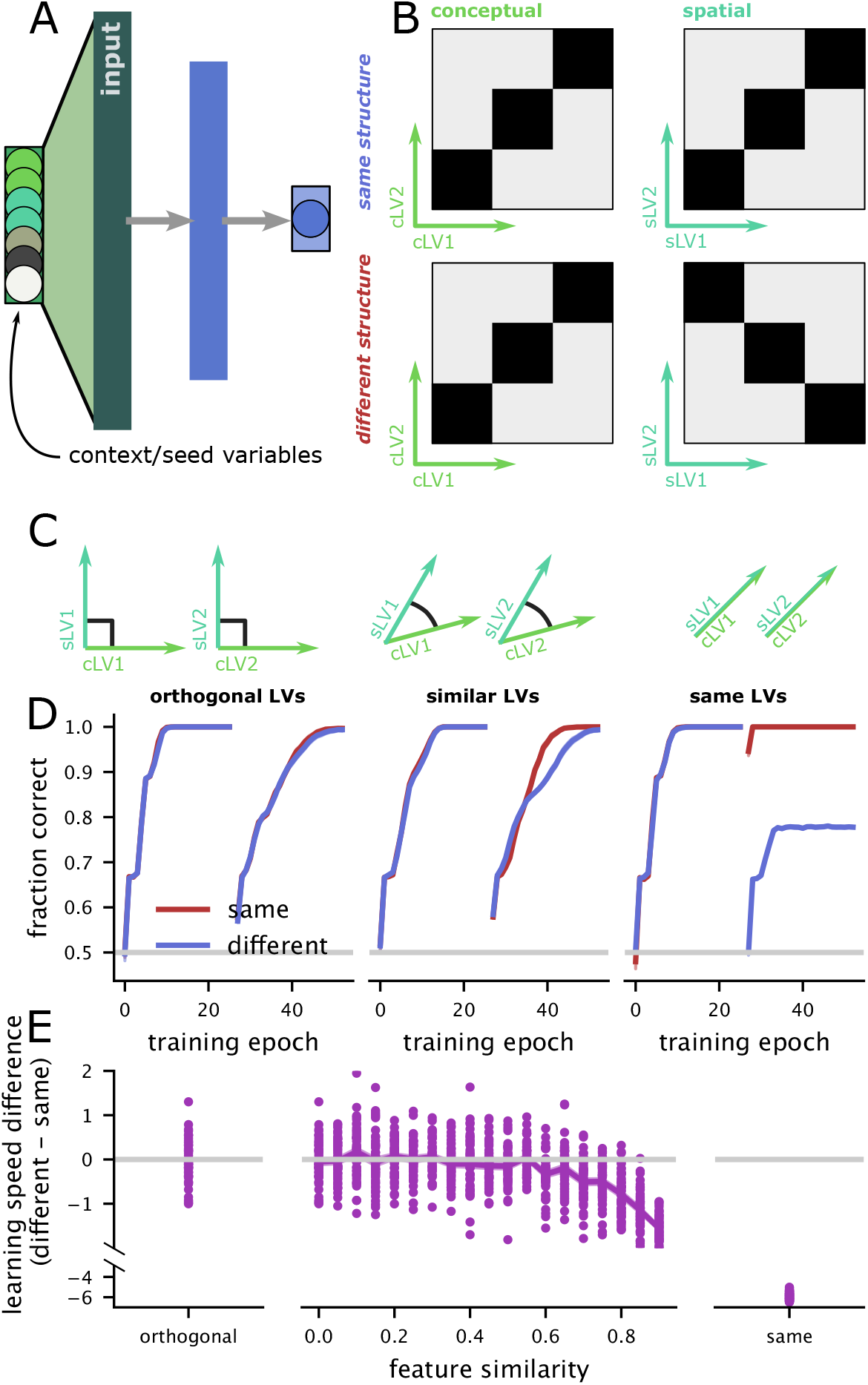
Blocked training in an RNN with aligned latent variables replicates our behavioral finding. **Panel A** Schematic of the neural network. There are two latent variables (LVs) relevant to the first context (cLV1 and cLV2), two relevant to the second (sLV1, sLV2), and two variables that signal the context (i.e., the seed identity in the experiment). There is a single hidden layer and one output unit. **Panel B** The tasks the network is trained to perform in two contexts (conceptual and spatial, left and right). The tasks are a coarser version of the experimental task and also have the same structure and different structure conditions. **Panel C** Schematic of the three input conditions for the network: (left) the latent variables are all orthogonal to each other; (middle) pairs of latent variables across the two contexts are encoded along semi-orthogonal directions; (right) there are only two, rather than four, total latent variables, and the latent variables across the different contexts are identical in the input to the network. **Panel D** The learning trajectories across 50 training epochs for each input structure, averaged over n = 50 random initializations of the model parameters. The error bars are the standard error of the mean (SEM), but are smaller than the line width. **Panel E** The average total difference in learning trajectory across the networks above for orthogonal inputs (left), a range of semi-orthogonal inputs (middle), and identical inputs (right). Each point shows a different initialization of the network.

### 3.1 Training Regime

Our results align with those of Flesch and colleagues (2018), showing that participants in the blocked regime perform better than those in the interleaved regime. This performance advantage persists into the testing phase. We found that the performance difference during testing is driven by improved memorization of old stimuli rather than by a benefit generalization to new stimuli. These results are also consistent with a contextual inference framework (Heald, Lengyel, & Wolpert, 2021, 2023; Shivkumar, Lengyel, & Wolpert, 2025), which predicts that blocked training facilitates the inference of task identity and thus supports the formation of distinct task representations. Interleaved training, by contrast, requires the learner to continuously infer the current task context on each trial, which may be particularly costly when tasks share structural features. While this pattern is consistent with previous work, our findings should be interpreted with caution, as the blocked and interleaved datasets were collected at different times and under slightly different experimental conditions. The literature on the benefits of different training regimes presents mixed evidence (Rohrer, Dedrick, & Stershic, 2015; Carvalho & Goldstone, 2014, 2015; Berens & Bird, 2022; Mi & Summerfield, 2026; Noh, Yan, Bjork, & Maddox, 2016). These discrepancies may stem from differences in what is being blocked or interleaved—specifically, whether the manipulation occurs at the level of entire tasks or within a single task at the level of individual stimulus configurations. In our study, tasks were blocked or interleaved, but it is possible that the effect would reverse if the manipulation were applied to the stimuli instead. That is, interleaving different stimulus configurations within a single task—rather than switching between different tasks—might encourage the integration of overlapping features and promote generalization in humans (see (Glitz, Liang, Barron, & Summerfield, 2026; Zhou, Singh, Tandoc, & Schapiro, 2023)). However, interleaving two tasks might create interference between representations that would otherwise be separated.

### 3.2 Transfer Learning in blocked training regime

Consistent with findings by Wu et al. (2020) and Lerousseau and Summerfield (2024), we observed a directional transfer effect in the blocked training regime, with performance on the second task influenced by the first. However, the direction of transfer in our study was reversed: whereas previous studies reported transfer from a spatial to a conceptual task, we found transfer from a conceptual to a spatial task. We suggest this discrepancy stems from differences in task nature. Wu et al.’s tasks involved navigation in spatial and conceptual spaces—that is, participants had to search for rewards by systematically exploring options based on two varying features, effectively navigating through a structured space of possibilities. These paradigms are likely grounded in spatial cognition. In such cases, spatial navigation may scaffold conceptual navigation, whereas spatial tasks may rely on more hardwired neural mechanisms hence remain unaffected by prior conceptual navigation. In contrast, our tasks involved classification, arguably more rooted in conceptual processing, as participants had to group stimuli based on abstract feature relationships rather than explore a physical or feature-based layout. This may have enabled knowledge acquired in the conceptual task to support learning in the spatial classification task, but not the reverse.

Lerousseau and Summerfield (2024) further showed that their spatial task promoted learning of a low-dimensional rule transferable to a conceptual context. In our case, learning the conceptual rule (e.g., “if light and water levels are similar, the seed grows”) required compressing a two-dimensional feature space into a one-dimensional rule space. Spatial classification, by contrast, may remain intrinsically two-dimensional, hence less generalizable. This asymmetry in representational complexity may account for the reversed direction of transfer observed in our study. One possible contributing factor is that acquiring an explicit rule structure may have provided a semantically rich task representation, thereby facilitating more stable task encoding and enabling more effective transfer of knowledge across tasks. This is consistent with evidence that semantically rich task labels and structured representations support robust task encoding and reduce interference during learning (Habibi, Verbeke, Senoussi, & Braem, 2025). An additional, non-mutually exclusive possibility is that this asymmetry is already present at the beginning of training, as suggested by a non-significant but numerically higher performance in the blocked condition in the conceptual task during the first two blocks (see Supplementary Figure 1), which may have provided an early advantage for conceptual-to-spatial transfer.

### 3.3 Cognitive control: Compositionality or Conjunctive Representation

Our findings also contribute to the cognitive control literature, which has long debated the nature of task representations, specifically whether they are compositional or conjunctive (Badre, 2024). Compositional representations are built from reusable components that transfer across tasks, enabling flexible behavior but at the cost of increased interference (Musslick & Cohen, 2019; Badre, Bhandari, Keglovits, & Kikumoto, 2021; Fusi, Miller, & Rigotti, 2016; Schwartenbeck et al., 2023; Bakermans, Warren, Whittington, & Behrens, 2025; Johnston, Fine, Yoo, Ebitz, & Hayden, 2024). In contrast, conjunctive representations bind task elements into task-specific codes, limiting generalizability but protecting from interference (Kikumoto & Mayr, 2020; Duncan, Doll, Daw, & Shohamy, 2018; Bhandari, Keglovits, & Badre, 2024). The pattern of interference we observed under interleaved training regime, particularly when tasks shared structural similarity, suggests that early in learning, representations may be compositional, allowing shared task features to overlap while being subject to interference. Over time, however, this interference diminishes, pointing toward a shift toward more conjunctive, task-specific representations. This learning-dependent transformation from compositional to conjunctive coding may reflect a mechanism by which the cognitive system balances generalization and interference as task demands and experience evolve (see also (Mill & Cole, 2025; Kikumoto & Mayr, 2020; O’Reilly & Rudy, 2001; Franklin & Frank, 2020; Boyle, Posani, Irfan, Siegelbaum, & Fusi, 2024)).

### 3.4 Consolidation theory and Cognitive Maps: Interference and Transfer

A similar dynamic is proposed in consolidation theories, which suggest that there is a critical window in which newly formed memories or knowledge are susceptible to interference from newly learned information (Lechner, Squire, & Byrne, 1999; Robertson, Pascual-Leone, & Miall, 2004; Zaki & Cai, 2024; Mutanen, Bracco, & Robertson, 2020). This period of instability allows the encoding of different knowledge to interact, potentially resulting in transfer and formation of high-level representations (Robertson, 2012; Mosha & Robertson, 2016; Robertson, 2018). Once knowledge is fully consolidated, transfer becomes less likely (Mosha & Robertson, 2016; Lu, Hummos, & Norman, 2024). Consolidation theories also predict that the benefits of interleaved training may become more pronounced after a more significant delay, as overnight consolidation could help stabilize the representations formed during interleaved practice. However, recent evidence suggests that the relative performance difference between blocked and interleaved training regimes does not change substantially over delays of 24 or 48 hours (Little & Nepangue, 2025; Habibi, Senoussi, Verbeke, & Braem, 2025). Our results align with and expand upon this framework. Our study suggests the existence of an additional layer of instability. Participants learning with an interleaved training regime experienced interference between tasks with similar structures, indicating the need for this knowledge to be stabilized. In contrast, participants in the blocked regime, who had more time for training on a single task, demonstrated knowledge transfer when the structures between tasks were similar. We propose a new relationship between the instability of newly learned information and transferability: high levels of instability of knowledge may be associated with interference, whereas some instability is necessary for knowledge transfer between tasks. On the other hand, full stabilization results in neither transfer nor interference (Mosha & Robertson, 2016). Furthermore, exploring longer delays may provide additional insight into the performance difference between the old and new configurations. Participants may have relied on both exemplar-based memory and rule-based knowledge during training. Because abstract rules may be retained better than exemplar-specific memories, delayed testing could reveal improved generalization relative to that observed here for novel configurations (Dominowski & Buyer, 2000; Elio & Anderson, 1984). Future studies will be needed to test this possibility directly. Finally, our findings provide additional evidence of the brain’s ability to abstract relationships between tasks in different domains into a higher-level representation. This representation is compatible with the concept of an amodal cognitive map—meaning a flexible, domain-general representation of structural relationships across various types of information, independent of the specific sensory or conceptual modalities involved (Behrens et al., 2018).

### 3.5 Conclusion

In conclusion, our study shows that the interaction between training regime and task structure similarity critically shapes representation learning. Participants trained under a blocked regime outperformed those in an interleaved regime during both training and testing. Crucially, when task structures were similar, the blocked regime facilitated knowledge transfer across tasks, though asymmetrically, with transfer occurring from the conceptual to the spatial task but not the reverse. In contrast, the same structural similarity impaired learning under the interleaved regime, where shared representations led to interference and reduced performance. This interference, however, decreased with practice, suggesting partial resolution over time. These findings underscore that similar task structures can either support (transfer) or hinder (interference) learning, depending on the training regime.

## 4 Methods

### 4.1 Participants

A total of 180 volunteers participated in the experiment (mean age = 28.9, SD = 4.4, 77 identified as males and 103 as females). The target sample size was determined based on two previous datasets using the same task paradigm that successfully identified differences between experimental groups (Menghi, Melega, Lidstrom, Renoult, & Penny, 2025; Menghi, Johnston, et al., 2025). All the participants were naive to the purpose of the experiment. 60 participants, belonging to the interleaved group condition, completed the experiment while we recorded MEG and their data were previously reported in (Menghi, Johnston, et al., 2025). The remaining 120 participants belonging to the blocked group condition completed the experiment in the behavioural lab. We discarded data from ten participants because their performance was below the chance level (56%) in both the spatial and the conceptual tasks. We performed analyses on the remaining 170 participants. At the end of the experiment, participants received reimbursement for their participation. All participants gave informed written consent, and the study procedure was approved by the local institutional review board of the University of Leipzig (Ethics Reference Number: 045/22-ek).

### 4.2 Apparatus and Stimuli

Participants completed the experiment inside a sound-attenuated, dimly lit, and magnetically shielded room. Stimuli were displayed on a monitor with a screen resolution of 1920 x 1080 pixels and a refresh rate of 60 Hz using the Psychophysics Toolbox (http://psychtoolbox.org/) (Brainard, 1997) for Matlab (Mathworks). Two images of seeds were used as cues to which kind of task (spatial or conceptual tasks) participants had to do. In the conceptual task, the opacity of a sun and a drop of water images governed the simulated amount of water and light received by the seed. Participants were instructed that greater opacity indicated higher levels of water or sunlight. Previous literature suggests that lower magnitudes are spatially represented toward the left and downward regions of space, whereas higher magnitudes are represented toward the right and upward regions (Viganò, Bayramova, Doeller, & Bottini, 2024; Loetscher, Bockisch, Nicholls, & Brugger, 2010; Hesse & Bremmer, 2017). In the spatial task, participants were presented with a black dot positioned within a white square, where the dot’s horizontal and vertical coordinates determined the virtual planting location of the seed. Stimuli were generated by sampling a subset of this two-dimensional spatial space.

#### 4.2.1 Task Structures

We used two different feature-outcome maps, which we refer to as structure (see Fig. 1). We used the same task structure that we used for our previous work (Menghi, Johnston, et al., 2025). These structures can be approximated by two different “diagonal” rules, allowing participants to learn the value of different spatial or conceptual combinations without sampling them first. Both task structures were defined using a deterministic mapping where the rounded value of the log-odds of the outcome was a quadratic function of stimulus characteristics, either the opacity of the two images or the position in space.

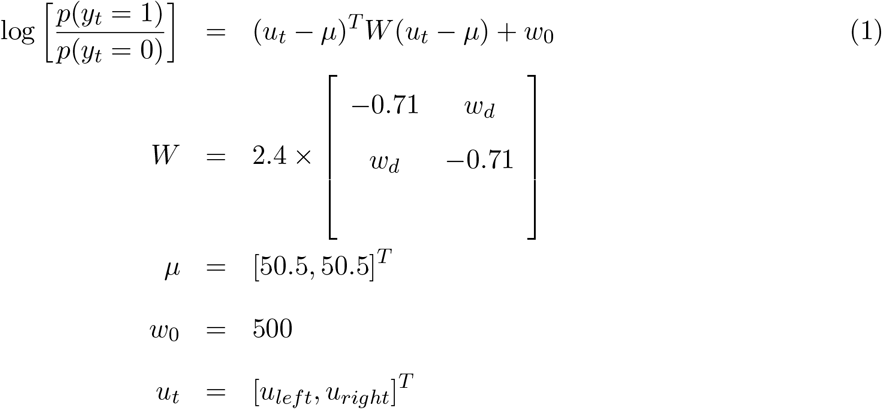

*u*_*t*_ here defines all the combinations of the feature of the task at hand. Water and light opacity for the conceptual task, and x and y positions for the spatial task. Flipping the sign of the wd parameter in this mapping produced the two structures depicted in Figure 1.

### 4.3 Experimental Design

Each participant was trained to do two tasks in either a blocked or an interleaved fashion. In the first task, the conceptual task, they had to learn the association between the opacity of a sun and a drop of water images and an outcome (grow or die); in the second one, they had to learn the association between a position in space and an outcome (grow or die). Importantly, spatial positions did not affect the conceptual task and conceptual features did not affect the spatial one. Participant trained in an interleaved fashion, during both training and test phases were presented with the two tasks interleaved. Participants trained with the blocked design were trained either with the spatial task first or with the conceptual one. Tasks were interleaved during the testing phase. Participants were randomly assigned to either a “Same structure” (SaSt) or a “Different structure” group (DiSt) and to either the blocked or the interleaved design. The features-outcome mappings for the SaSt group were generated using *wd* = 0.71 for both spatial and conceptual tasks (See previous paragraph and equation 1). The features-outcome mappings for the DiSt group were generated using *wd* = 0.71 for the conceptual task and *wd* = −0.71 for the spatial task.

### 4.4 Procedure

The experiment comprises four distinct phases, as illustrated in the C panel of Figure 1 and lasted about 2 hours. The overall experimental procedure and task environment were adapted from our previous work (Menghi, Johnston, et al., 2025). Throughout the training and test phases, participants completed a computerized task in which they took on the role of scientists working in a biology laboratory. In this simulated environment, the laboratory had cultivated two novel plant species, and participants were required to learn the environmental conditions that determined the successful germination of the corresponding seeds.

#### 4.4.1 Training

As illustrated on the right part of Fig. 1, during the first phase, the training task, in each trial participants will first be prompted with one of two seeds cueing which task to focus on and then the stimuli appeared and stayed on screen for 2500 ms maximum or until a response was made. Responses were made on a standard button box, one button indicating a prediction of ‘grow’ and another predicting “die”. Responses not given within the required time constitute “missed trials”. Right after the button press feedback was provided, saying “correct” if the prediction was correct, “incorrect” if it was not and “too slow” if they missed the trial (no response within 2500 ms). By making predictions and receiving feedback, participants learned the association between relevant features and the outcome “grow” or “die” as in a classic associative learning task. During the training phase, participants were exposed to 18 different configurations in both the spatial and conceptual tasks. Each configuration was presented 8 times, with each repetition paired with a different irrelevant configuration from the other task dimension. For example, in the spatial condition, each spatial configuration was paired with 8 different conceptual configurations (4 associated with “grow” and 4 associated with “die”). This resulted in a total of 288 training trials (18 configurations × 8 repetitions × 2 tasks). Importantly, some stimuli (i.e., coordinates within the two-dimensional feature spaces) overlapped across the two contexts, whereas others were unique to each context.

#### 4.4.2 Test

During the second phase, the test tasks, we tested participants’ knowledge about the structures learned during the training. No feedback was provided during the test phase. We presented old and new stimuli to test for differences in memorization, transfer and generalization. Some of the new stimuli were new in both contexts and some were previously encountered (old) in one context but entirely new in the other context. Importantly, performance did not statistically differ between these two types of novel stimuli in either the blocked group (old in one context/new in the other: *t*(114) = 0.114, *p* = 0.909; new in both contexts: *t*(114) = 0.488, *p* = 0.625) or the interleaved group (old in one context/new in the other: *t*(114) = −1.685, *p* = 0.097; new in both contexts: *t*(114) = 0.458, *p* = 0.648). Therefore, these two types of novel stimuli were collapsed in the main analyses. During the testing phase, participants were exposed to 28 different configurations in both spatial and conceptual tasks. Like during the training session, each configuration was repeated 8 times, with a different irrelevant configuration. For example, in the spatial condition, each spatial configuration was paired with 8 conceptual configurations (4 associated with “grow” and 4 with “die”). The total number of trials in the testing phase was thus 896 (28 configurations × 8 repetitions x 2 tasks). This phase is divided into “first testing session” and “second testing session” with 8 repetition each, separated by a navigation task that will be analysed separately.

#### 4.4.3 Navigation

The navigation task addresses a distinct question that will be analyzed separately. Here we provide a short description of the task. Participants were presented with one of the two seeds and an expected outcome: “grow” or “die”. Participants were then asked to navigate the appropriate (spatial or conceptual) space, by varying the two features (x and y for the spatial seed, amount of sun and rain for the conceptual seed), locate a position on the map and “plant” the seed according to the outcome requested. The task lasted approximately 15 minutes and was administered between the first and second testing sessions. Importantly, performance did not differ significantly before and after the navigation task in three of the four experimental conditions (Blocked–Same: *t*(58) = 1.574, *p* = 0.121; Blocked–Different: *t*(56) = 0.930, *p* = 0.356; Interleaved–Same: *t*(26) = 1.244, *p* = 0.225), with only a small decrease observed in the Interleaved–Different condition (*t*(26) = 2.084, *p* = 0.047; pre-test *M* = 0.730, post-test *M* = 0.718). Given the minimal differences observed, the two testing sessions were combined in the main analyses.

## Data and Code Availability

The code for the neural network was written in python, using TensorFlow and numpy. The code is freely available on https://github.com/wj2/modularity/tree/master. Behavioural data and analyses implemented in Matlab software are available from https://github.com/Nich0Me/TrainingRegimeRepositor

## Acknowledgments

We thank Yvonne Wolf-Rosier, Anna Krauspe and Dareen Darwish for their invaluable assistance with participant recruitment and data collection, Kerstin Schumer for her exceptional management of the project. C.F.D.’s research is supported by the Max Planck Society, the European Research Council (ERC-CoG GEOCOG 724836), the Kavli Foundation, the Jebsen Foundation, the Centre of Excellence scheme of the Research Council of Norway—Centre for Neural Computation (223262/F50), The Egil and Pauline Braathen and Fred Kavli Centre for Cortical Microcircuits, and the National Infrastructure scheme of the Research Council of Norway—NORBRAIN (197467/F50).

## Supplementary Materials

**Supplementary Figure 1:**
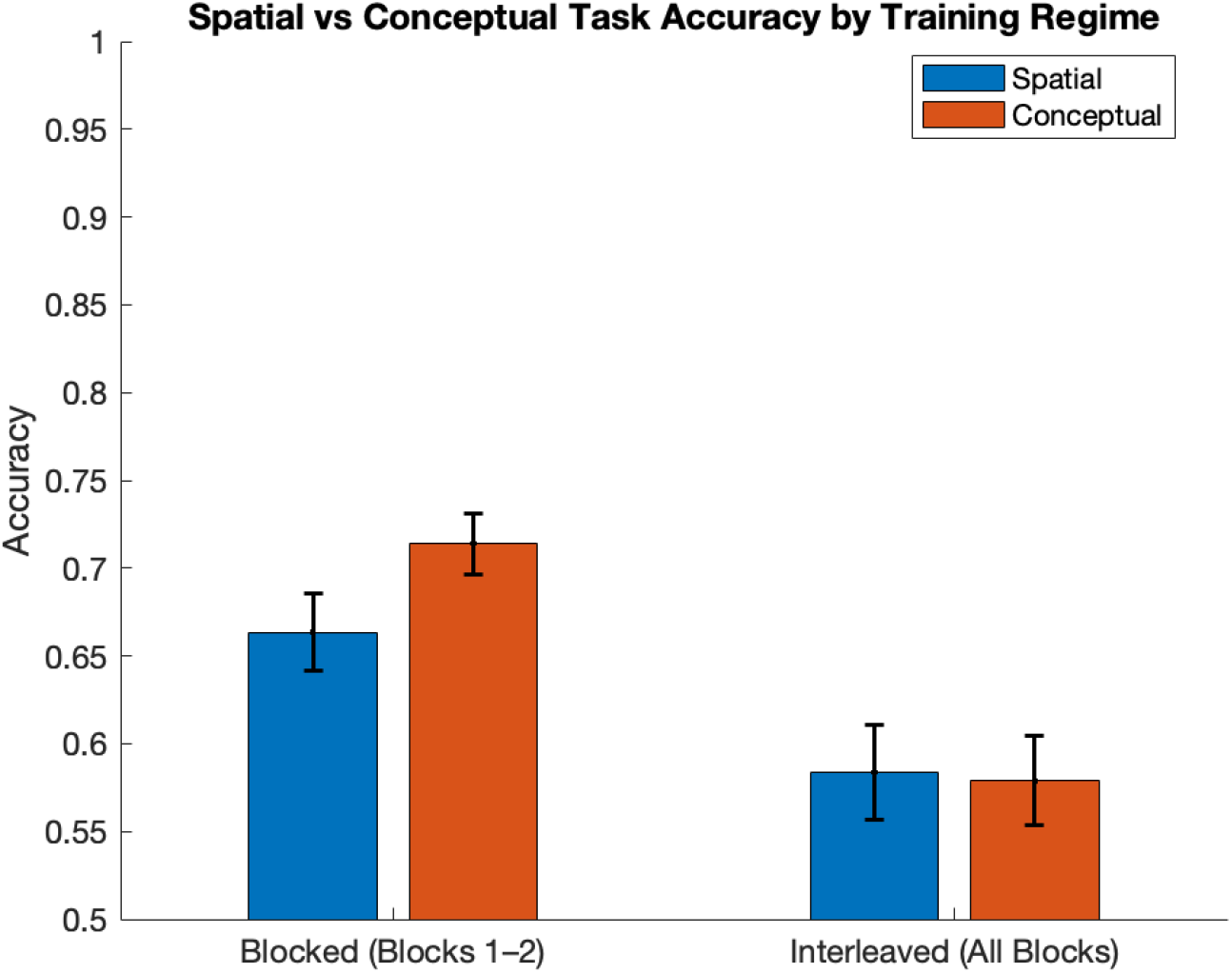
Task performance across spatial and conceptual tasks in blocked and interleaved regimes. On the left: mean accuracy (± SEM) for the blocked condition, shown separately for participants starting with the spatial versus conceptual task. A between-subjects comparison indicated a non-significant trend towards higher performance in the conceptual task relative to the spatial task (t(117) = 1.804, p = 0.074). Right: mean accuracy (± SEM) for the interleaved condition, shown for spatial and conceptual tasks within participants. No reliable difference was observed between task types in the interleaved regime (t(58) = 0.385, p = 0.702).

**Supplementary Figure 2:**
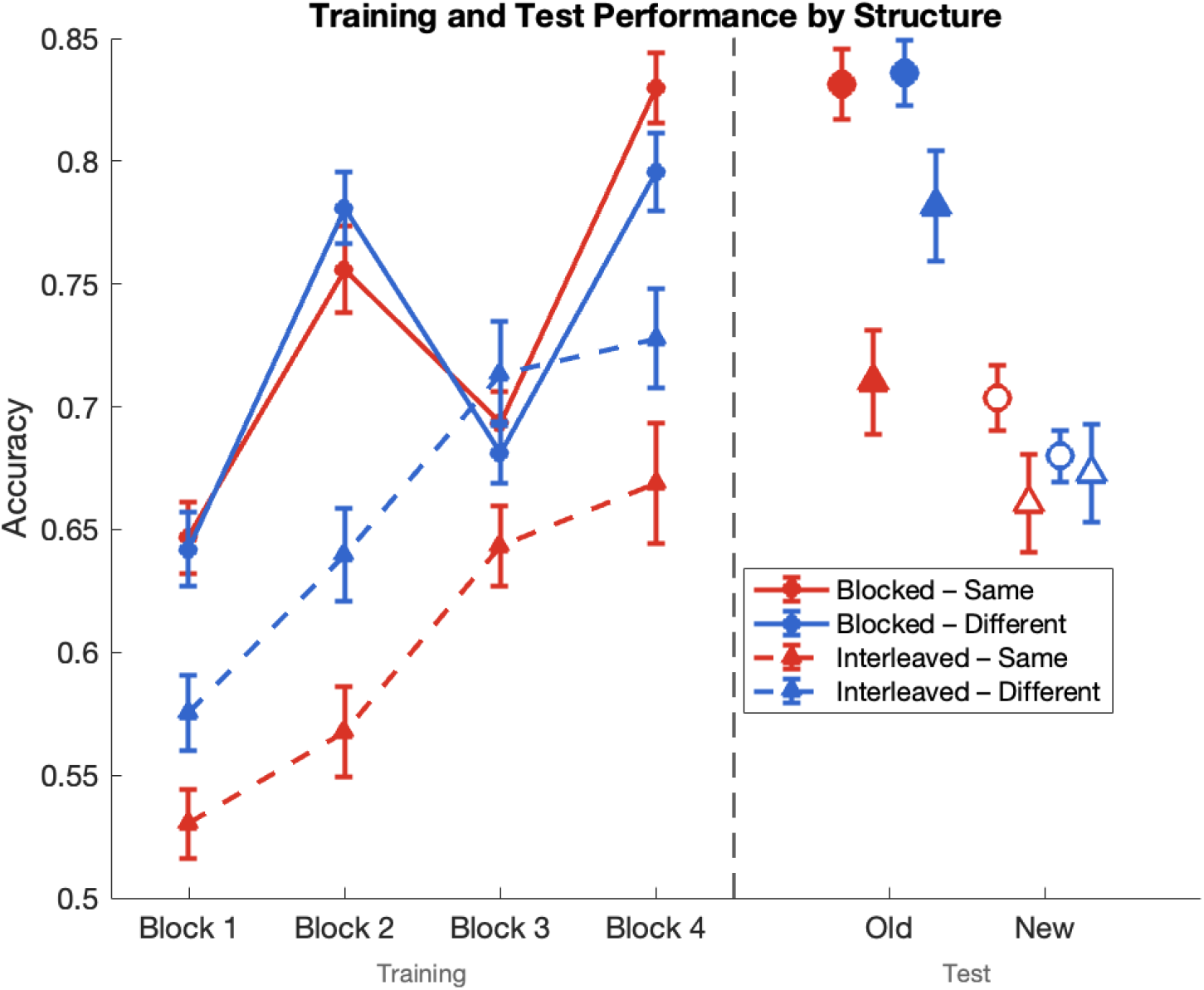
Training and test accuracy by training regime, task structure and novelty. Mean accuracy (± SEM) is shown across four training blocks (left of dashed line) and at test for old and new items (right of dashed line) for four conditions: Blocked–Same (red solid line, circles), Blocked–Different (blue solid line, circles), Interleaved–Same (red dashed line, triangles), and Interleaved–Different (blue dashed line, triangles). Filled markers indicate old items; open markers indicate new items. During training, blocked conditions began at higher accuracy (0.65) and showed a dip where the new task was starting, reaching 0.83 (Same) and 0.80 (Different) by Block 4. Interleaved conditions started lower (0.53–0.58) and improved steadily throughout training, reaching 0.67–0.73 by Block 4. At test, old item accuracy was statistically indistinguishable from the last training block in the Blocked–Same condition (t(58) = 0.15, p = .884), but exceeded it in all other conditions (Blocked–Different: t(56) = 3.96, p < .001; Interleaved–Same: t(26) = 2.68, p = .013; Interleaved–Different: t(26) = 3.25, p = .003), suggesting learning effect. New item accuracy was significantly lower than Block 4 in blocked conditions (Same: t(58) = 9.10, p < .001; Different: t(56) = 7.84, p < .001) and in the Interleaved–Different condition (t(26) = 2.63, p = .014), but did not differ from Block 4 in the Interleaved–Same condition (t(26) = 0.34, p = .734).

